# Physical principles of phase-separation action on chromatin looping associated to pathogenic gene activation

**DOI:** 10.1101/2025.05.20.654944

**Authors:** Sougata Guha, Andrea Fontana, Alex Abraham, Simona Bianco, Andrea Esposito, Mattia Conte, Sumanta Kundu, Ciro Di Carluccio, Francesca Vercellone, Florinda Di Pierno, Mario Nicodemi, Andrea M. Chiariello

## Abstract

Phase-separation of chimeric proteins resulting from genetic mutations has been shown to trigger aberrant chromatin looping, contributing to disease development, including cancer. However, the physical mechanisms regulating these processes are not yet fully understood. In this study, we employ polymer physics models of chromatin and numerical simulations to investigate the relationship between phase-separation of proteins and chromatin structure. We demonstrate that a simple model, including only protein-protein and protein-chromatin interactions, effectively explains the aberrant looping observed around oncogenes, such as *PBX3*, in cells expressing the NUP98-HOXA9 chimeric protein, which is associated with leukemia. In this scenario, looping occurs through a switch-like mechanism controlled by the concentration of the chimera and its affinity with chromatin. Moreover, when incorporating the presence of extruding factors in a more complex model, similar results are observed, indicating a mild dependence of this looping mechanism on loop-extrusion activity. Finally, leveraging our numerical simulations, we propose potential strategies to inhibit the formation of enhancer-gene loops by directly targeting the chimeric protein with interfering molecules.

## Introduction

The spatial organization of genomes within the nucleus of a cell is fundamental to its gene regulation and cell function^1^. A growing amount of experimental evidence is highlighting phase-separation (PS) of protein condensates as one of the key physical mechanisms controlling chromatin organization and gene regulation. In combination with processes such as loop-extrusion^2,3^, polymer micro-phase separation has been discovered to be an organization mechanism of chromatin 3D conformation, from the scale of TADs^4–6^, to of A/B compartments^7^, which partition the genome in regions associated with transcriptionally active and repressed chromatin, respectively^1,8,9^.

The assembly of protein aggregates, leads to the formation of stable, membraneless organelles which are implicated, e.g., in mediating long-range interactions between genes and regulators^10,11^, in forming larger condensates, such as Cajal bodies^12,13^, in the formation of complex multiphase aggregates made of different molecular species (as nucleolar inter-chromosomal hubs^14,15^ or highly specific enhancer hubs, e.g., in olfactory sensory neurons^16^) and in the enhancement of gene bursting^17^.

The formation of phase-separated aggregates is driven by the tendency of some proteins to interact with themselves, typically through their intrinsically disordered regions (IDRs)^12^. Relevant examples of proteins capable of such homo-typic interactions include mediator complex and RNA Polymerase II (RNA PolII)^18–20^, coactivators^10^ and transcription factors (TFs)^11,21,22^. In addition, condensates can be formed by multiple molecules mediated by hetero-typic interactions, such as between proteins and RNA in the nucleolus^12,15,23^.

On the other hand, phase-separation mechanisms can be associated with various diseases^24,25^, including cancer and neurodegenerative disorders. For example, liquid-liquid phase-separation of the RNA-binding protein hnRNPA1, a protein related to severe diseases as amyotrophic lateral sclerosis and proteinopathy, has been shown to drive pathogenic fibrillization^26^. Another relevant example is aberrant phase-separation of the protein HMGB1 caused by frameshift variants, which leads to nucleolar disfunction and is linked to rare genetic diseases, as malformation syndromes^27^. In this case, mutations can alter the dynamics and the properties of protein condensates, leading to the formation of pathological aggregates that disrupt normal cellular functions. Additionally, polymer models based on phase separation have been used to study chromatin misfolding in congenital diseases^28,29^. In cancer, chimeric proteins resulting from genetic mutations, such as the NUP98-HOXA9 fusion found in leukemia^30,31^, can drive aberrant chromatin looping and altered gene expression through phase-separation^32,33^.

Here, we investigate with polymer models and Molecular Dynamics (MD) simulations the physical mechanisms underlying the formation of phase-separated protein clusters and their interaction with chromatin. Using as case study the above mentioned NUP98-HOXA9 chimera, we show that a simple model of the chimeric protein, exhibiting the typical hallmarks of phase-separation, and a chromatin filament with binding sites for the chimera representing a genomic locus around the *PBX3* gene, naturally defined using ChIP-seq data, is sufficient to explain aberrant loop formation and is validated against Hi-C data. In agreement with experiments^32^, inhibition of phase-separation properties of the chimera, e.g., by switching off its self-interaction, disrupts such contacts. Furthermore, similar results are found also around the *MAP2K5* gene locus, in which CTCF and cohesin play a role in organizing chromatin structure, likely through loop-extrusion mechanism. Indeed, the model correctly predicts the formation of gene-enhancer loops upon phase-separation of the chimeric protein as well as loop loss when phase-separation only is inhibited. Interestingly, loop-extrusion is mildly affected as the model indicates new loops from a slightly increased processivity for the extruders when phase-separation is lost. Analysis of 3D trajectories reveals that chimeric proteins bound to chromatin exhibit reduced mobility, in agreement with experimental data^32^. Finally, we investigate possible strategies to inhibit the formation of aberrant looping by introducing in the system interfering molecules. Our simulations suggest that aberrant looping is efficiently reduced by targeting the DNA-binding region of the chimeric protein (i.e., the HOXA9 domain), which interestingly is a known target for specific inhibitors and central in pharmaceutical research^34^.

Overall, our study investigates the physical principles underlying the interplay between phase-separation and chromatin looping, employing polymer models applied to NUP98-HOXA9 chimera as case study, and show that a simple model including protein-protein and protein-chromatin interactions, with and without loop-extrusion, quantitatively explains the aberrant looping observed in Hi-C data^32^. Given the generality of these models, our results could be extended to any biological system with similar features providing a general computational framework able to quantitatively explore chromatin aberrant looping, even in pathogenic contexts, in a highly controlled manner.

## Results

We study the role of the NUP98-HOXA9 chimeric protein as a chromatin organizer and the interplay between its phase-separation tendency and its ability to induce aberrant pathogenic looping. To this aim, we consider recently published Hi-C data^32^ in human 293FT cells expressing NUP98-HOXA9 in wild-type (WT) condition and in human 293FT cells expressing a mutated version of NUP98-HOXA9, in which a phenylalanine-to-serine (FS-mutated) substitution in their IDRs impairs the formation of phase-separated condensates.

### Minimal modelling of NUP98-HOXA9 recaps features of protein coarsening and dynamics

The chimeric protein NUP98-HOXA9 results from a chromosomal translocation, i.e., t(7;11)(p15;p15), typically associated with leukemia^30,31^, leading to the fusion of a segment of the nucleoporin-encoding *NUP98* gene, containing intrinsically disordered regions (IDRs) generally prone to drive phase-separation^10,11,35^, and a region of the *HOXA9* gene, which encodes for a DNA-binding factor^30,32^. Importantly, liquid-liquid phase-separation of NUP98-HOXA9 chimeric protein has been shown to drive chromatin looping and, likely, potentiates gene activation^32^. This system is therefore an excellent case study to investigate, through computational modelling, the interplay between phase-separation and alteration of chromatin structure, with consequent impact on gene regulation^36^.

To do so, we first studied the properties of a protein model including as key ingredients self-interaction and affinity with binding sites located on the chromatin filament^37^ (Methods). We considered as model for NUP98-HOXA9 chimera a molecule including a region which can interact with chromatin (i.e., HOXA9 domain) and a region capable of self-interaction (i.e., NUP98 IDRs domain, Fig. 1a, Methods). As dictated by polymer physics, Molecular Dynamics simulations show formation of molecular clusters when self-interaction affinity E_IDR_ and concentration ρ_IDR_, which control system thermodynamic state, are above threshold (Fig. 1b, Methods). Also, cluster formation occurs through coalescence events (Fig. 1c), i.e., first small droplets are formed and then they fuse, as observed in microscopy experiments^32^. At equilibrium, statistics of clusters depend on system control parameters as, e.g., higher concentrations return fewer and more abundant clusters (Fig. 1d, Supplementary Figs. 1a,b).

**Figure 1.**
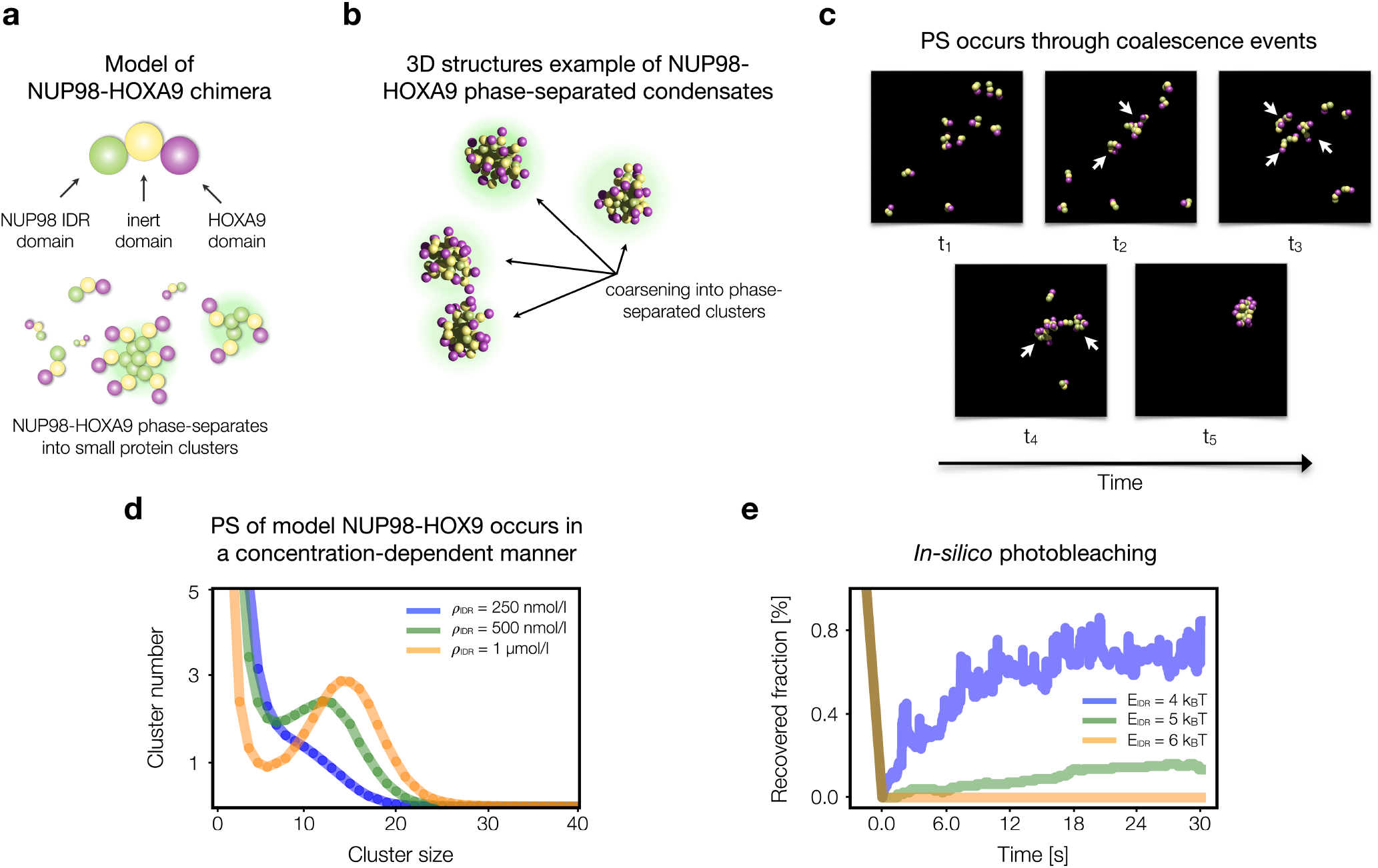
Minimal model of NUP98-HOXA9 chimera recaps phase-separation hallmarks. **a:** Model of NUP98-HOXA9 chimeric protein. HOXA9 domain is colored in purple, the IDR of NUP98 domain in green. IDR attractively self-interact among themselves. **b:** 3D snapshots from MD simulation of NUP98-HOXA9 protein clusters. **c:** Protein cluster dynamics evolves through coalescence events. White arrows denote single proteins coarsening together. **d:** Number of clusters at different size depends on protein concentration. **e:** Average fraction of proteins, within clusters, recovered from external environment as a function of time, at different affinities E_IDR_, shows that phase-separation (PS) properties of model NUP98-HOXA9 condensates depend on affinity.

To understand if such a model for NUP98-HOXA9 protein is also able to recap different phase-separation regimes (e.g., liquid-like and solid-like), we studied the dynamics of the clusters and calculated the time interval required to exchange proteins with the surrounding environment, by tracking in time position of proteins belonging to a fixed cluster (Supplementary Fig. 1c, Methods). Essentially, we simulated an *in-silico* version of fluorescence recovery after photobleaching (FRAP) experiment, which is commonly used to investigate the nature of phase-separated clusters^10–12^. As expected, the estimated recovery curve depends on affinity E_IDR_ (Fig. 1e), as lower affinities exhibit faster recovery times, indicating a higher turn-over of proteins within the aggregate, whereas higher affinities required longer time to alternate proteins. Interestingly, this latter case could be tagged as solid-like condensate (very stable and limited molecule exchange with external environment) and has been recently indicated as condensation process for Lhx2/Ebf1 transcription factors in olfactory sensory neurons^16^. Of note, for low affinity values (i.e., E_IDR_ around 4 k_B_T) we estimate time scales to reach plateau around 15-20 seconds (Methods), compatible with results for liquid-liquid phase-separation (LLPS) driven by IDR interactions reported in literature^10,11,35^. Analogous considerations hold for the value of the plateau of recovered proteins^10^, which is in the range 60-80% for affinities around 4 k_B_T, and much lower (10-20%) for higher affinities (above 5 k_B_T). Taken together, those results confirm that our model of NUP98-HOXA9 is very general and exhibits the typical hallmarks of LLPS, as shown in experiments^32^.

### Polymer models describe aberrant looping around PBX3 oncogene locus

Next, we tested the above-described model of NUP98-HOXA9 as chromatin organizer and the interplay between its phase-separation tendency and looping induction on chromatin (Fig. 2a). To this aim, we studied the architecture of a real genomic locus and considered a 600 kb long region (chr9: 128,300,000-128,900,000 bp, hg19 assembly) around the oncogene *PBX3*, which is up-regulated and involved in leukemias (especially acute myeloid leukemia, AML)^38–40^. Hi-C data in 293FT cells expressing NUP98-HOXA9 (ref.^32^) reveal the presence of strong loops involving the *PBX3* gene (Fig. 2b, upper panel, data from ref.^32^). The loops anchor points correspond to peaks of NUP98-HOXA9 ChIP-seq data^32^ and also H3K27ac, indicating the presence of regulatory elements involved in the loops, also reported in public enhancers databases^41^ (Fig. 2b, lower panels). Importantly, no apparent CTCF sites are involved in the 3D organization of this locus, suggesting that those loops are formed in a CTCF-independent manner^32^.

**Figure 2.**
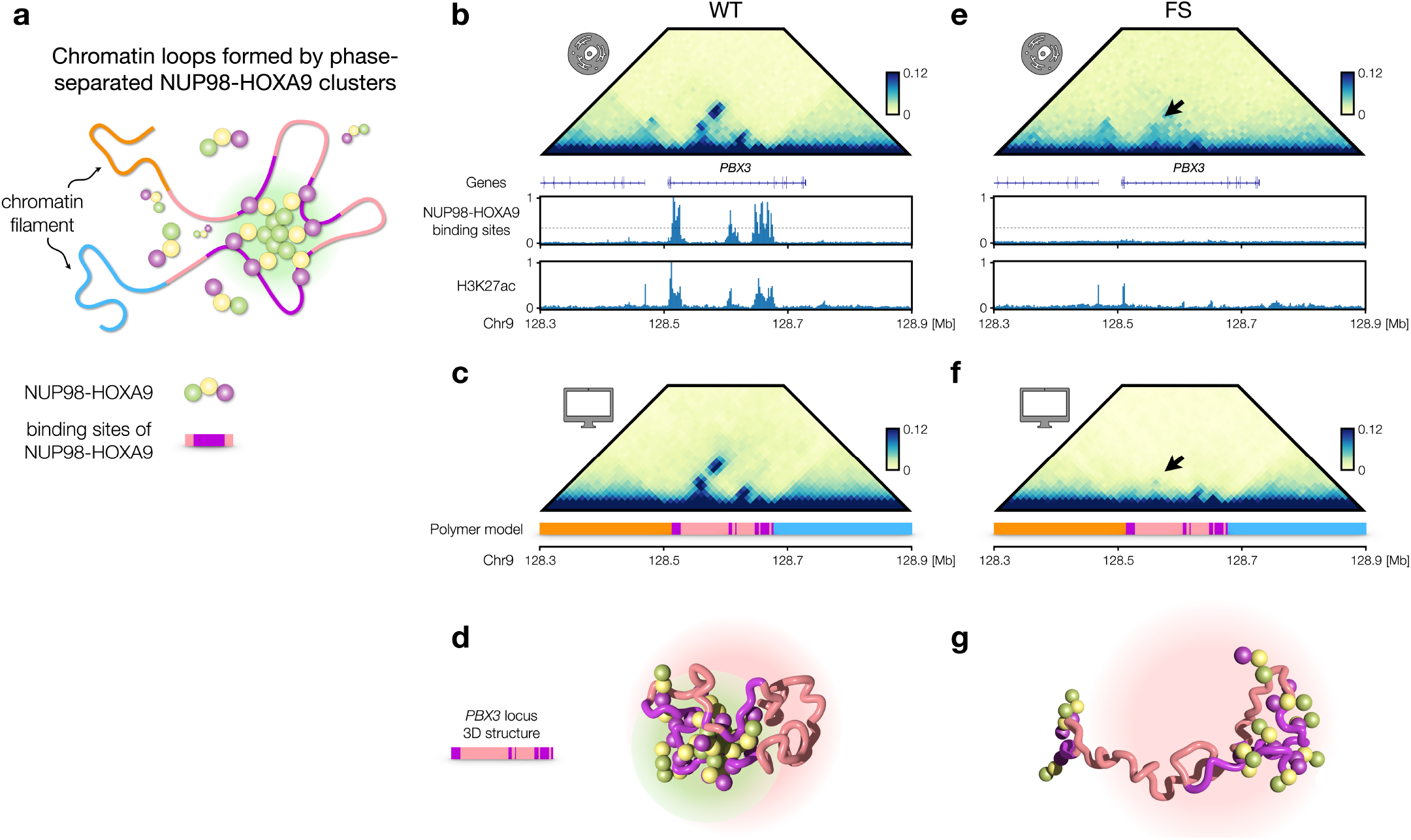
Polymer physics and protein phase-separation describe aberrant looping at PBX3 locus. **a:** Schematic of polymer model including chromatin filament and NUP98-HOXA9 chimeric proteins. Different colors highlight different polymer regions. Binding sites for NUP98-HOXA9 are represented in purple. **b:** Hi-C data of the genomic region chr9: 128.3-128.9 Mb (hg19 assembly) in 293FT cells (data taken from ref.^32^), expressing wild-type (WT) NUP98-HOXA9 chimeric proteins. Below, ChIP-seq signal of NUP98-HOXA9 chimera (used to define specific binding sites) and H3K27ac. **c:** *In-silico* contact map obtained from MD simulations of WT NUP98-HOXA9 protein-chromatin system accurately captures aberrant looping. Distance-corrected correlation with experimental Hi-C is r’ = 0.65. Below, color scheme of the polymer model is shown. **d:** Example of 3D structure (around the *PBX3* gene) taken from an MD simulation of WT model, including polymer and chimeras. **e:** Hi-C data of the genomic region chr9: 128.3-128.9 Mb (hg19 assembly) in 293FT cells (data taken from ref.^32^), expressing phenylalanine-to-serine (FS) mutated NUP98-HOXA9 chimeric proteins. Black arrow denotes loss of aberrant looping. **f:** *In-silico* contact map obtained from MD simulations of FS-mutated NUP98-HOXA9 protein-chromatin system predicts loss of aberrant looping (black arrow). **g:** As **d**, for the FS-mutated model.

So, we defined a simple polymer model with binding sites for NUP98-HOXA9 whose location is determined by using ChIP-seq data (Fig. 2c, lower panels, Methods), as done in other similar polymer models to describe chromatin organization^6,20,42,43^. Then, by means of Molecular Dynamics simulations (Methods) we generated ensembles of 3D structures by varying the system parameters (concentration ρ_IDR_ and affinity E_IDR_, keeping fixed the affinity with chromatin E_HOXA9_ for sake of simplicity). Loop formation occurs through switch-like behavior, as highlighted by the sharp decrease of polymer size in the region containing the *PBX3* gene (Supplementary Fig. 2a, left panel) when E_IDR_ is above threshold, indicating that the loops are triggered by the phase-transition of the protein cluster^36^. Intriguingly, hypothesizing that E_IDR_ is chemically related to the number of phenylalanine and glycine (FG) aminoacidic repeats in the IDRs of NUP98, this is in agreement with the loss of phase-separation experimentally observed, by microscopy and immunoblotting^32^, for engineered NUP98-HOXA9 proteins with a reduced number of FG repeats in the IDR region and with the reduced binding of NUP98-HOXA9 to *PBX3* gene (measured by ChIP-qPCR) when the same mutated chimera with less FG repeats is expressed^32^. Analogous results are found when ρ_IDR_ is increased, although the polymer size reduces more gradually (Supplementary Fig. 2a, right panel). By aggregating different polymer structures (Methods) we are able to reconstruct *in-silico* the contact map of the locus, which is highly correlated to Hi-C data after the phase-transition (Fig. 2c, upper panel), with the characteristic strong loops (distance-corrected Pearson r’ = 0.65, Methods). Snapshot of 3D structures shows in space the 3D folding of *PBX3* gene and the protein cluster mediating the contacts between the gene and its regulatory elements (Fig. 2d). Next, we tested whether the model is able to predict loss of looping upon inhibition of phase-separation. To this aim, we compared experimental Hi-C maps of *PBX3* gene with NUP98-HOXA9 chimera in which IDRs have been mutated through a phenylalanine-to-serine mutation (i.e., FS-mutated)^32^ to disable formation of phase-separated droplets. Here, Hi-C data shows the complete loss of aberrant looping (Fig. 2e, upper panel). In our model, inhibition of phase-separation is easily implemented by just switching-off self-interaction of NUP98 IDR part of the molecule (i.e. setting E_IDR_ = 0, Methods), with ρ_IDR_ and E_HOXA9_ parameters kept unchanged. In agreement with Hi-C data, simulated contact map shows the complete loss of loops (Fig. 2f, upper panel), although the interaction between HOXA9 with chromatin is still enabled and some residual contacts persist. 3D snapshot of the system (Fig. 2g) in this condition highlights the loss of loops in the chromatin structure. Therefore, the model correctly predicts with accuracy that CTCF-independent aberrant looping is driven by phase-separation of NUP98-HOXA9 chimera.

### Interplay between loop-extrusion and aberrant looping driven by phase-separation

We then investigated a genomic locus with aberrant looping in which CTCF and cohesin play a role in organizing chromatin structure (Fig. 3a). To this aim, we considered as case study a genomic region centered around the *MAP2K5* gene (chr15: 67,550,000-68,350,000 bp, hg19 assembly). Although *MAP2K5* is not directly related with leukemias, it is associated to other pathogenic contexts, e.g., neurological and movement disorders^44^, metabolic disorders^45^ and its pathway is also dysregulated in cancers^46,47^. Hi-C data at this locus exhibit strong loops around the gene corresponding to NUP98-HOXA9 ChIP-seq peaks (Fig. 3b, upper panel). Furthermore, strong CTCF peaks surround NUP98-HOXA9 peaks, likely indicating the presence of loop-extrusion activity (Fig. 3b, lower panels). Therefore, we considered a polymer model equipped with binding sites for NUP98-HOXA9 defined using ChIP-seq data (Methods), as previously described, and anchor points for loop-extruding factors (typically identified as cohesin molecules^2,48^), defined from most prominent CTCF ChIP-seq peaks^32^ (Methods). Extruding factors actively slide along the polymer to simulate loop-extrusion^3,49,50^. Hence, loop-extrusion, phase-separation of NUP98-HOXA9 proteins and chromatin-NUP98-HOXA9 interactions simultaneously occur in the system and cooperatively determine chromatin organization at this locus (Methods). Again, by performing MD simulations, we were able to generate populations of 3D structures and, correspondingly, *in-silico* contact maps which accurately recapitulates the features (i.e. loops and boundaries) shown in Hi-C data (distance-corrected Pearson correlation r’ = 0.69, Fig. 3c). Again, loop formation is associated to a sharp decrease in polymer local size upon increase of control parameters, E_IDR_ and ρ_IDR_, above threshold (Supplementary Fig. 2b). Snapshot of 3D structures shows in space the 3D folding of *MAP2K5* gene and the protein cluster mediating the contacts between the gene and its regulatory elements (Fig. 3d).

**Figure 3.**
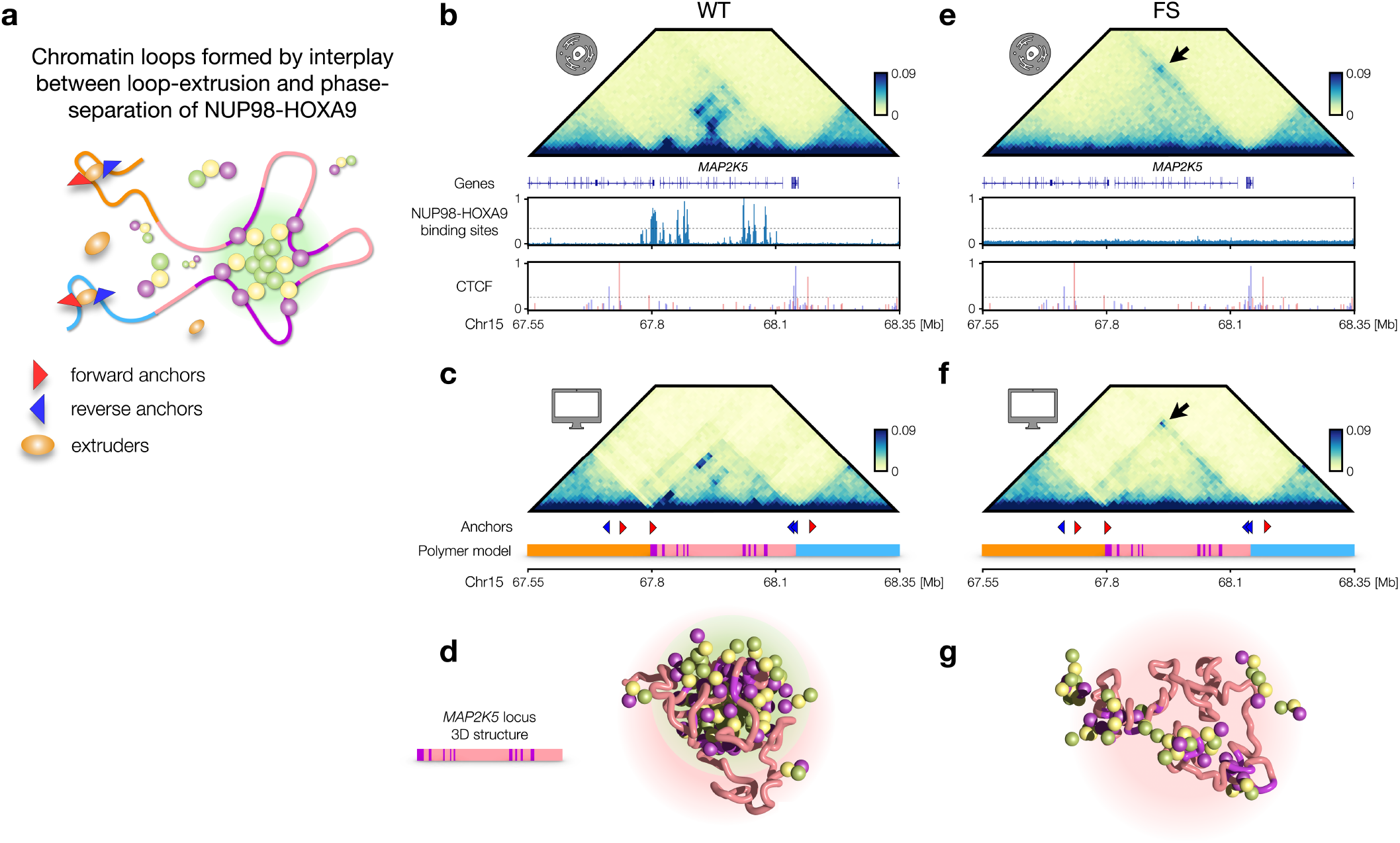
Interplay between loop-extrusion and aberrant looping driven by phase-separation. **a:** Schematic of polymer model including chromatin filament and NUP98-HOXA9 chimeric proteins. Different colors highlight different polymer regions. Binding sites for NUP98-HOXA9 are represented in purple. Arrowheads indicate forward (blue) and reverse (red) anchor points on the chromatin filament for extruding factors, shown in orange. **b:** Hi-C data of the genomic region chr15: 67.55-68.35 Mb (hg19 assembly) in 293FT cells (data taken from ref.^32^), expressing WT NUP98-HOXA9 chimeric proteins. Below, ChIP-seq signal of NUP98-HOXA9 chimera (used to define specific binding sites) and anchor probability taken from CTCF ChIP-seq signal (Methods) to simulate loop-extrusion. **c:** *In-silico* contact map obtained from MD simulations of WT NUP98-HOXA9 protein-chromatin system accurately captures aberrant looping. Distance-corrected correlation with experimental Hi-C is r’ = 0.69. Below, color scheme of the polymer model and location of anchor points (with their orientation) are shown. **d:** Example of 3D structure (around the *MAP2K5* gene) taken from an MD simulation of WT model, including polymer and chimeras. **e:** Hi-C data of the genomic region chr15: 67.55-68.35 Mb (hg19 assembly) in 293FT cells (data taken from ref.^32^), expressing FS-mutated NUP98-HOXA9 chimeric proteins. Black arrow denotes loss of aberrant looping. **f:** *In-silico* contact map obtained from MD simulations of FS-mutated NUP98-HOXA9 protein-chromatin system predicts loss of aberrant looping (black arrow). Note the enhanced loop deriving from enhanced processivity in loop-extrusion. **g:** As **d**, for the FS-mutated model.

Next, we investigated the system with inhibition of phase-separation of NUP98-HOXA9 chimera, as described above. Hi-C data in 293FT cells expressing NUP98-HOXA9 FS-mutated chimera show the loss of strong loops associated to the NUP98-HOXA9 peaks and a new loop detected in the FS-mutated condition, not present in the WT condition (Fig. 3e and Supplementary Fig. 2d, upper panel)^32^. As before, we simulated inhibition of phase-separation by setting to E_IDR_ = 0, keeping unchanged the other model parameters (i.e., E_HOXA9_, ρ_IDR_, number of extruders and loop-extrusion processivity, Methods). In agreement with Hi-C data, loops mediated by NUP98-HOXA9 phase-separation are lost and the TAD organization is mainly driven by loop-extrusion (Supplementary Fig. 2c), with a CTCF mediated loop mildly formed with respect to the WT one (Supplementary Fig. 2d, lower left panel). Importantly, by increasing the loop-extrusion processivity (approximately by 40%, Methods) we were also able to recover the presence of the loop observed in the Hi-C data of FS-mutated (Fig. 3e) and its intensity (Supplementary Fig. 2d, lower right panel), suggesting that dissolution of the phase-separated cluster impacts loop-extrusion activity by enhancing extruder processivity or, conversely, formation of phase-separated clusters constrains loop-extrusion activity with consequent reduction of the path length of cohesin along the chromatin fiber. 3D snapshot of the system (Fig. 3g) in this condition highlights the loss of loops in the chromatin structure.

### Phase-separation affects protein mobility

Next, we studied dynamic properties of the chromatin-protein system to quantitatively test the impact of chromatin occupancy by NUP98-HOXA9 on the protein mobility. To this aim we investigated the mobility of the NUP98-HOXA9 chimera under different physical conditions (Fig. 4a). Specifically, we first computed the three-dimensional linear displacement (Methods) of single NUP98-HOXA9 proteins in WT and FS-mutated conditions in the model of *PBX3* locus. We find that, at equilibrium, in WT condition displacements tend to fluctuate around average values significantly lower than the FS condition (Fig. 4b). The result is confirmed when the single-molecule *in-silico* experiment is repeated on independent simulations (Fig. 4c, one-sided t-test p-value = 10^−22^). Importantly, this result is in agreement with single molecule live imaging experiments^32^ performed on human 293FT cells expressing WT and FS-mutated NUP98-HOXA9 chimera and indicates a more constrained dynamics of the proteins due to the presence of phase-separated cluster^36^ and chromatin-binding^51,52^.

**Figure 4.**
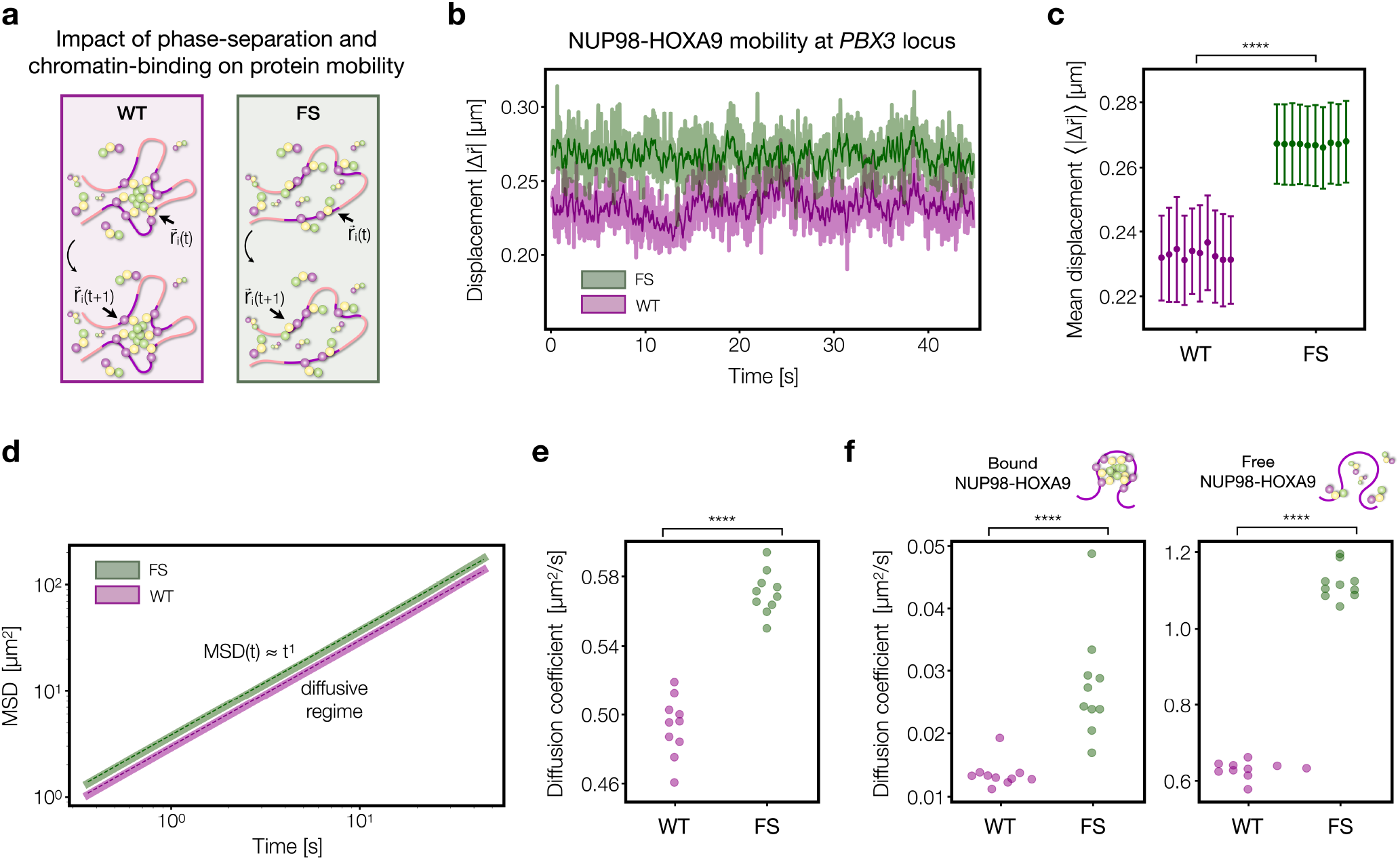
Phase-separation and chromatin-binding influence protein mobility. **a:** NUP98-HOXA9 mobility is investigated in both WT and FS conditions. **b:** Displacements of single proteins in WT (purple) and FS (green) during time. **c:** Distributions of mean values of single protein displacements over independent simulations of polymer model of *PBX3* locus in WT and FS conditions are statistically different (one-sided t-test p-value = 10^−22^). **d:** Mean square displacement (MSD) of chimeras exhibits power law with time compatible with exponent 1 (dashed lines), in both WT and FS conditions. **e:** Distributions of average diffusion coefficients over independent simulations in WT and FS conditions are statistically different (one-sided t-test p-value = 10^−11^). **f:** Estimates of diffusion coefficients in bound and free states (Methods), in both WT and FS conditions. The estimated numerical range is comparable with experimental values^32^.

Analogous results are found for the system with the polymer model of *MAP2K5*, although with a less marked difference between WT and FS-mutated conditions (Supplementary Figs. 3a,b, one-sided t-test p-value = 10^−21^), suggesting a role for loop-extrusion in enhancing molecule mobility, likely due extrusion activity, in agreement with recent results from polymer models based on loop-extrusion^53,54^. This result highlights also how loop-extrusion activity can impact protein mobility, although no direct interaction exists between extruders and proteins as extrusion acts on chromatin fiber only. Analysis of mean square displacement (MSD, Methods) in WT and FS-mutated conditions (Fig. 4d) returned a similar scenario, with scaling exponent α ~ 1 and estimated diffusion coefficients in WT significantly lower than FS condition (Fig. 4e, one-sided t-test p-value = 10^−11^). In line with the previously described locus, diffusion coefficients estimated from the model of *MAP2K5* locus on large time scales in WT tend to be lower than the FS condition (Supplementary Figs. 3c,d, one-sided t-test p-value = 10^−9^), suggesting that the observed reduction of mobility is a general behavior of proteins induced by chromatin occupancy.

As more stringent test for our model, we made estimates of diffusion coefficients considering proteins in a bound-to-chromatin or freely-diffusing state (two-state kinetic model^55,56^, Methods) and found values in the approximate range 0.5-1.5 μm^2^/s for free and ≪ 0.1 μm^2^/s for bound condition (Fig. 4f and Supplementary Fig. 3e), in agreement with the above mentioned live imaging experimental data^32^.

Taken together, those results shows that the model predicts an impact on protein mobility due to phase-separation and chromatin-binding which is confirmed by experimental imaging data of NUP98-HOXA9 proteins from entire cell nuclei^32^.

### Interference mechanisms of PS and possible strategies to inhibit aberrant looping

The formation of phase-separated NUP98-HOXA9 aggregates mediating loops between genes and regulators, as in the case of the *PBX3* locus, and the concomitant up-regulation for several other genes shown in ref.^32^, suggest a possible causal relation between phase-separation and gene aberrant activation in leukemia and indicate this process as putative, general mechanism driving gene mis-regulation in other pathogenic contexts^32^. Therefore, we investigated, leveraging on our *in-silico* models, possible strategies to interfere with formation of aberrant contacts. As case study, we focused again on the above-described model of *PBX3* locus. Specifically, we tested the effect of adding in the system a new molecule (Fig. 5a) which can interact only with NUP98-HOXA9 chimera so to interfere with its looping capacity. In other words, this simple system simulates the action of a hypothetical drug having NUP98-HOXA9 as molecular target. Specifically, we considered as target region the HOXA9 domain, which is a typical target for some inhibitors^34^, e.g., HXR9, DB1055 (ref.^57^), DB818 (ref.^58^), HTL-001 (ref.^59^) and has a central role for pharmacological research in leukemia^60^.

**Figure 5.**
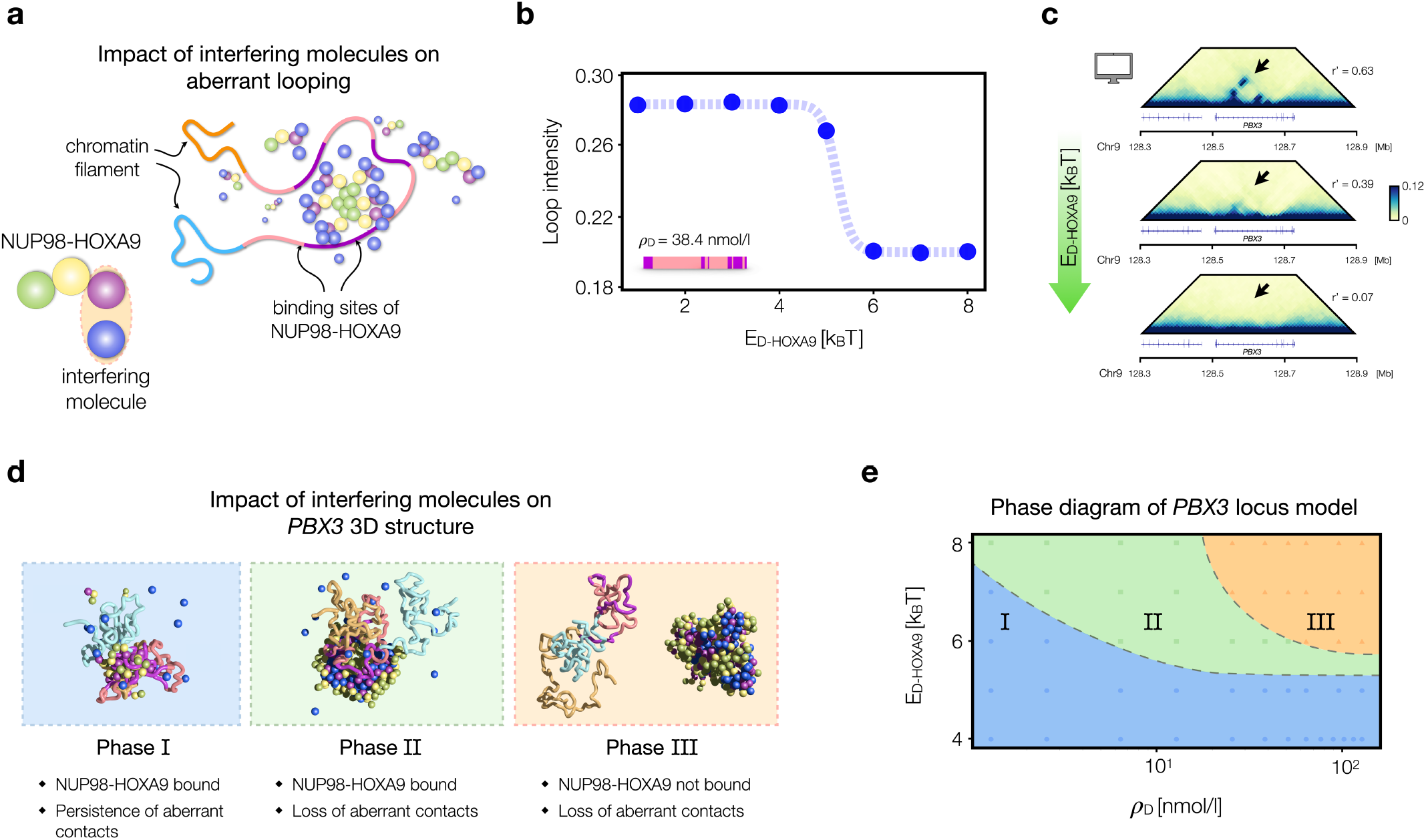
Interference mechanisms of PS and possible strategies to inhibit aberrant looping. **a:**Inhibition of aberrant looping is investigated by introducing new molecules in the system (in blue) capable of attractive interaction with the NUP98-HOXA9 chimera. **b:** Aberrant loop intensity in the *PBX3* locus polymer model depends on the affinity E_D-HOXA9_ between interfering molecules and HOXA9 part of the chimera (for a fixed concentration ρ_D_) and drastically drops after a threshold value. A sigmoid is also shown for visualization purposes. **c:** *In-silico* contact maps of the *PBX3* locus for different affinities E_D-HOXA9_ of interfering molecules. Black arrows denote progressive loss of aberrant looping. **d:** 3D configurations for low (left panel), intermediate (central panel) and high (right panel) affinities E_D-HOXA9_. **e:** Phase diagram of the system including NUP98-HOXA9 chimeras, polymer of *PBX3* locus and interfering molecules (Methods). In the blue region, i.e., low E_D-HOXA9_ affinities and ρ_D_ concentrations, no loss of aberrant contacts is observed (Phase I); in green (i.e., intermediate E_D-HOXA9_ and ρ_D_, Phase II) and orange (i.e., high E_D-HOXA9_ and ρ_D_, Phase III) regions, loss of aberrant contacts is detected.

Then, to evaluate the performances of the interfering molecule, we simulated the system by varying its binding affinity E_D-HOXA9_ and its concentration ρ_D_ (Methods) and monitored the formation of the aberrant loops (Figs. 5b-d, Methods). We find that loops disappear by increasing E_D-HOXA9_ above a threshold value (Fig. 5b), for a fixed concentration ρ_D_. Visual inspection of 3D snapshots for different values of E_D-HOXA9_ (Fig. 5d) offered the opportunity to physically interpret the loss of contact. Interestingly, two possible cases emerge. For moderately high values of E_D-HOXA9_, aberrant contacts are lost because a bigger, phase-separated cluster made of NUP98-HOXA9 and interfering molecules still interacts with chromatin but pushes apart the gene from the regulatory elements (Fig. 5d, central panel), in a chromatin restructuring mechanism driven by phase-separation, analogous to the mechanically induced chromatin re-modelling through the CasDrop experimental method^61^. For higher affinities E_D-HOXA9_, the molecules completely interfere and the NUP98-HOXA9 cluster detaches from chromatin, with consequent loss of contacts (Fig. 5d, right panel). Analogous results are found when concentration ρ_D_ is varied, keeping fixed E_D-HOXA9_ (Supplementary Figs. 4a-c), indicating this behavior as a general, controllable feature of this system.

To support this conclusion, we performed extensive simulations varying E_D-HOXA9_ and ρ_D_ as control parameters and calculated the size of the region enriched with binding sites for NUP98-HOXA9 chimera and its distance from the cluster of proteins and interfering molecules (Methods). Both curves exhibit approximately sigmoidal shapes upon increase of E_D-HOXA9_ (Supplementary Fig. 4d) or ρ_D_, with clearly different transition values, highlighting the different mechanisms behind the observed loss of contacts. Then, by classifying the equilibrium states, we built the phase diagram of the system. We were able to identify three main regions defining the state of the system based on the presence of the aberrant contact and on the mechanisms leading to its loss (Fig. 5e). To check the robustness of these results, we verified that the same mechanisms occur also in the *MAP2K5* model (Supplementary Figs. 5a-e).

Finally, we performed additional simulations by targeting the NUP98 domain of the chimeric protein and found that in the considered parameter range no relevant loop loss is detected (Supplementary Fig. 5f), indicating targeting of HOXA9 as more efficient strategy to inhibit the aberrant contact.

In synthesis, those results highlight how the modelling approach can be useful to discover strategies to inhibit pathogenic mechanism at a molecular level (e.g., targeting chromatin-binding HOXA9 region of the NUP98-HOXA9 chimera) and to optimize them, e.g., identifying transition thresholds.

## Discussion

In this work, we investigated the interplay between protein phase-separation and chromatin looping, with a particular focus on consequent pathogenic gene activation. To this aim, we considered a computational model of NUP98-HOXA9, a chimeric protein resulting from a leukemia-associated translocation, in which phase-separation of proteins is coupled with polymers simulating real genomic regions, by integrating ChIP-seq data of CTCF and NUP98-HOXA9 binding sites from human 293T cells expressing NUP98-HOXA9. Then, by performing Molecular Dynamics simulations we generated ensembles of 3D structures to explore genome architecture under different physical conditions. It emerges that a simple model with protein-protein and chromatin-protein interactions is able to quantitatively explain CTCF-independent aberrant loops observed in experimental Hi-C data, such as at the genomic locus around the *PBX3* oncogene, and it is able to predict a switch-like loss of aberrant looping upon inhibition of phase-separation of NUP98-HOXA9 chimera. Furthermore, incorporation of loop-extrusion in the polymer model allows to describe loci where CTCF plays a role in organizing chromatin structure and suggests an impact of phase-separation on extruder processivity along the chromatin fiber. Finally, we show that targeting of HOXA9 domain in the chimera with interfering molecules turns out to be a robust and effective strategy to inhibit aberrant looping, in line with experimental research which identified inhibitors for HOXA9 (ref.^34^).

Collectively, those results elucidate the role of phase-separation mechanism on chromatin looping and highlight the validity of computational models to investigate those systems in a highly controlled way. Therefore, they can be useful for a wide range of applications including, but not limited to: supporting experimental approaches to find and optimize research of inhibitors for proteins capable of phase-separation and involved in pathogenic pathways, e.g., BET bromodomains^62^; studying the transcriptome regulation through condensates of proteins in wild-type or mutated forms^63^; complementary validation of computational predictions from approaches combining experiments and deep learning (e.g., ProtoGPS model^64,65^) to predict the physicochemical properties of biomolecular condensates. As looping through phase-separation could be a general mechanism involved in different diseases, it could be also an important tool to new pharmacological design^25^.

## Acknowledgements

We acknowledge funding from Fondazione Compagnia di San Paolo - Linea PoC Launchpad and from INFN, project “COBRA “COmputer Based RNA Analysis and prediction” nell’ambito dell’avviso pubblico relativo al bando a cascata, D.R. n. 750 del 28/02/2024 dell’Università degli Studi di Bari, per lo spoke 7 “BIOCOMPUTING” del progetto PNRR CN00000041 “National Center for Gene Therapy and Drugs based on RNA Technology”, Missione 4, Componente 2, Investimento 1.4, finanziato dall’Unione europea - NextGenerationEU - CUP H93C22000430007”.

## Author contributions

AMC and MN designed and coordinated the project. SG and AF run computer simulations and performed data analysis with support from AA. AMC, MN, AF and SG wrote the manuscript with input from all the authors.

